# Structure-Based Neural Network Protein-Carbohydrate Interaction Predictions at the Residue Level

**DOI:** 10.1101/2023.03.14.531382

**Authors:** Samuel W. Canner, Sudhanshu Shanker, Jeffrey J. Gray

## Abstract

Carbohydrates dynamically and transiently interact with proteins for cell-cell recognition, cellular differentiation, immune response, and many other cellular processes. Despite the molecular importance of these interactions, there are currently few reliable computational tools to predict potential carbohydrate binding sites on any given protein. Here, we present two deep learning models named CArbohydrate-Protein interaction Site IdentiFier (CAPSIF) that predict carbohydrate binding sites on proteins: (1) a 3D-UNet voxel-based neural network model (CAPSIF:V) and (2) an equivariant graph neural network model (CAPSIF:G). While both models outperform previous surrogate methods used for carbohydrate binding site prediction, CAPSIF:V performs better than CAPSIF:G, achieving test Dice scores of 0.597 and 0.543 and test set Matthews correlation coefficients (MCCs) of 0.599 and 0.538, respectively. We further tested CAPSIF:V on AlphaFold2-predicted protein structures. CAPSIF:V performed equivalently on both experimentally determined structures and AlphaFold2 predicted structures. Finally, we demonstrate how CAPSIF models can be used in conjunction with local glycan-docking protocols, such as GlycanDock, to predict bound protein-carbohydrate structures.

## 1 Introduction

The carbohydrate-protein handshake is the first step of many pathological and physiological processes (1, 2). Pathogens attach to host cells after their lectins successfully bind to surface carbohydrates (or glycans) (3–6). The innate and adaptive immune systems utilize carbohydrate signatures present on cellular and subcellular surfaces to recognize and destroy foreign components (7, 8). Glycosaminoglycans (GAGs) bind to membrane proteins of adjacent cells for cell-cell adhesion and to regulate intracellular processes (9–11). Despite the biological importance of these carbohydrate-protein interactions, there are few carbohydrate-specific tools leveraging the vast Protein DataBank (PDB) and recent advances in machine learning (ML) to elucidate the binding of carbohydrates at a residue level.

Knowledge of carbohydrate-protein interactions has been leveraged to develop therapeutic candidates to neutralize infections and inspire proper health function (6, 12). One bottleneck in designing carbohydrate-mimetic drugs is obtaining residue-level interaction knowledge through methods such as structural data and/or mutational scanning profiles (12–14). Further, in some studies, computational tools have been used to predict docked structures, refine bound carbohydrates, or extract dynamic information (14–16).

Recent developments in deep learning (DL) have substantially enhanced the theoretical modeling of proteins and protein-protein interactions. For example, neural networks can design stable proteins with unique folds using graph representations (17, 18). 3D structures can be predicted with programs such as IgFold and Alphafold2 (AF2) (19, 20). Predicted 3D atomic coordinates can be probed to determine ligand or protein binding capabilities using neural networks such as Kalasanty or dMaSIF (21, 22).

Recent computational studies have demonstrated new ways to explore protein-carbohydrate interactions. Our lab has also contributed to the advancement of this field by adding the following, (1) a shotgun scanning glycomutagenesis protocol to predict the stability and activity of protein glycovariants (23), and (2) the GlycanDock algorithm to refine protein-glycoligand bound structures (24).

Recently there have been developments in small molecule binding site predictors. Small molecule binding site predictors typically fall into four categories: template, geometry, energy, or machine learning based (25). Template based strategies, such as 3DLigandSite (26), search datasets for sequence and/or structurally related ligand binding proteins to assess prospective binding sites. Geometry based methods, like FPocket (27), search the surface of proteins for pockets and cavities. Energy based methods, such as FTMap (28), use probe molecules to scan the surface of a protein to determine the energetic favorability of binding. Recently, machine learning techniques, such as Kalasanty (21), have emerged and outperformed previous classical site prediction algorithms, commonly with convolutions on a 3D voxel grid containing atomistic information (29, 30).

Although there are many general small molecule binding site predictors (21, 28, 31), few tailored algorithms exist for prediction of protein-carbohydrate sites. In 2000, Taroni *et al*. performed an analysis of carbohydrate binding spots using the solvation potential, residue propensity, hydrophobicity, planarity, protrusion, and relative accessible surface area to construct a function to predict carbohydrate binding sites (32). In 2007, Malik and Ahmad created a neural network to predict the carbohydrate binding sites using their constructed Procarb40 dataset, a collection of 40 proteins, with leave one out validation (33). In 2009, Kulharia built InCa-SiteFinder to predict carbohydrate and inositol binding sites by leveraging a grid to construct an energy-based method for predicting binding sites (34). Tsai *et al*. constructed carbohydrate binding probability density maps using an encoding of 30 protein atom types as an input to a machine learning algorithm (35). Later, Zhou, Yang and colleagues developed two methods to predict carbohydrate binding sites, (1) a template-based approach named SPOT-Struc (36) and (2) a support vector machine (SVM) named SPRINT-CBH that leverages sequence-based features (37). Tsia (35) and SPOT-Struc (36) have achieved Matthews correlation coefficients (MMCs) of 0.45 on test sets of 108 and 14 proteins, respectively. The increased size of the protein databank and the improvements in deep learning methods now presents an opportunity to train and test more broadly.

Larger protein-carbohydrate structural databases now include UniLectin3D (38) and ProCaff (39). UniLectin3D focuses on lectins bound to carbohydrates, containing 2406 structures; however, it contains many redundant structures and is currently limited to 592 unique sequences. ProCaff lists 552 carbohydrate-binding protein structures and their binding affinities under various conditions; however, many structures are only available in the unbound form.

Many drug targets, from pathogen-lectins to aberrant selectins, are carbohydrate binding proteins (6, 13, 40). Understanding the physiological response and determining a glycomimetic drug to neutralize the infection requires residue-level knowledge (40). Currently, DL algorithms LectinOracle (41) and GlyNet (42) predict lectin-carbohydrate binding on a protein level; however, pharmaceutical development requires residue-level information.

In this work, we develop two DL methods for residue-level carbohydrate-binding site prediction. The two methods have different architectures, one using voxel convolutions and one using graph convolutions. We also present a dataset of 808 bound nonhomologous protein chain-carbohydrate structures and use it to train and test both models. We compare the performance of the models with each other and with FTMap (28) and Kalasanty (21). Then, we evaluate the performance of the models on AlphaFold2 (20) predicted versus experimentally determined structures. Finally, we present a proof-of-concept pipeline to predict bound protein-carbohydrate structures.

## 2 Results

### 2.1 Dataset for carbohydrate-protein structures

To construct a method to predict carbohydrate-protein interactions, we needed a large and reliable dataset to use for training and testing. The dataset should contain as many non-homologous structures as possible to avoid biasing to specific folds. By filtering the PDB (43), we constructed a dataset of 808 high accuracy (< 3 Å resolution), nonhomologous (30% sequence identity), and physiologically relevant experimental structures (by manually removing buffers), spanning 16 carbohydrate monomer species. In these structures, 5.2% of the protein residues contact carbohydrates **(Supplementary File S1)**. The final dataset consists of 808 structures, which we split into 521 training structures, 125 validation structures, and 162 test structures.

### 2.2 CAPSIF uses deep neural networks to predict carbohydrate interaction sites

We constructed convolutional neural networks (CNNs) named CArbohydrate-Protein Site IdentiFier (CAPSIF) to predict carbohydrate binding residues from a protein structure. CNNs were initially developed for images, exploiting the spatial relationship of nearby pixels for prediction tasks. They have been applied to predict protein structure (44–46) and small molecule binding pockets of proteins (21). To predict carbohydrate binding residues using structural information, we created two CAPSIF CNN architectures, CAPSIF:Voxel (CAPSIF:V) and CAPSIF:Graph (CAPSIF:G).

Since a protein can change its side chain conformations upon binding a small molecule or carbohydrate (from *apo* to *holo*), we sought a protein representation that is robust to these and other binding induced changes. We chose a residue-level representation, using only the Cβ positions of all residues (or Cα in glycine), since the Cβ position is frequently equivalent in both the *apo* and *holo* states (47). Both CAPSIF architectures use the following features: unbound solvent accessible surface area (SASA) of each residue, a backbone orientation (architecture specific), and encodings of amino acid properties, including hydrophobicity index (0 to 1) (48), “aromatophilicity” index (0 to 1) (49), hydrogen bond donor capability (0,1), and hydrogen bond acceptor capability (0,1) (**Methods**/**Table 3**).

The first CAPSIF architecture, CAPSIF:V, is a 3D voxelized approach to learn carbohydrate binding pockets. CAPSIF:V uses a UNet architecture, which comprises a grid with a series of convolutions compressing and then decompressing the data to its original size with residual connections to previous layers of the same size. For each grid, we used an 8 Å^3^ voxel size where CAPSIF:V encodes each residue’s β carbon (Cβ) into a corresponding voxel. CAPSIF:V predicts a label *P*(carbohydrate-binding residue) for each voxel on the initial grid (**Figure 1A; Methods/Figure 6)**.

**Figure 1:**
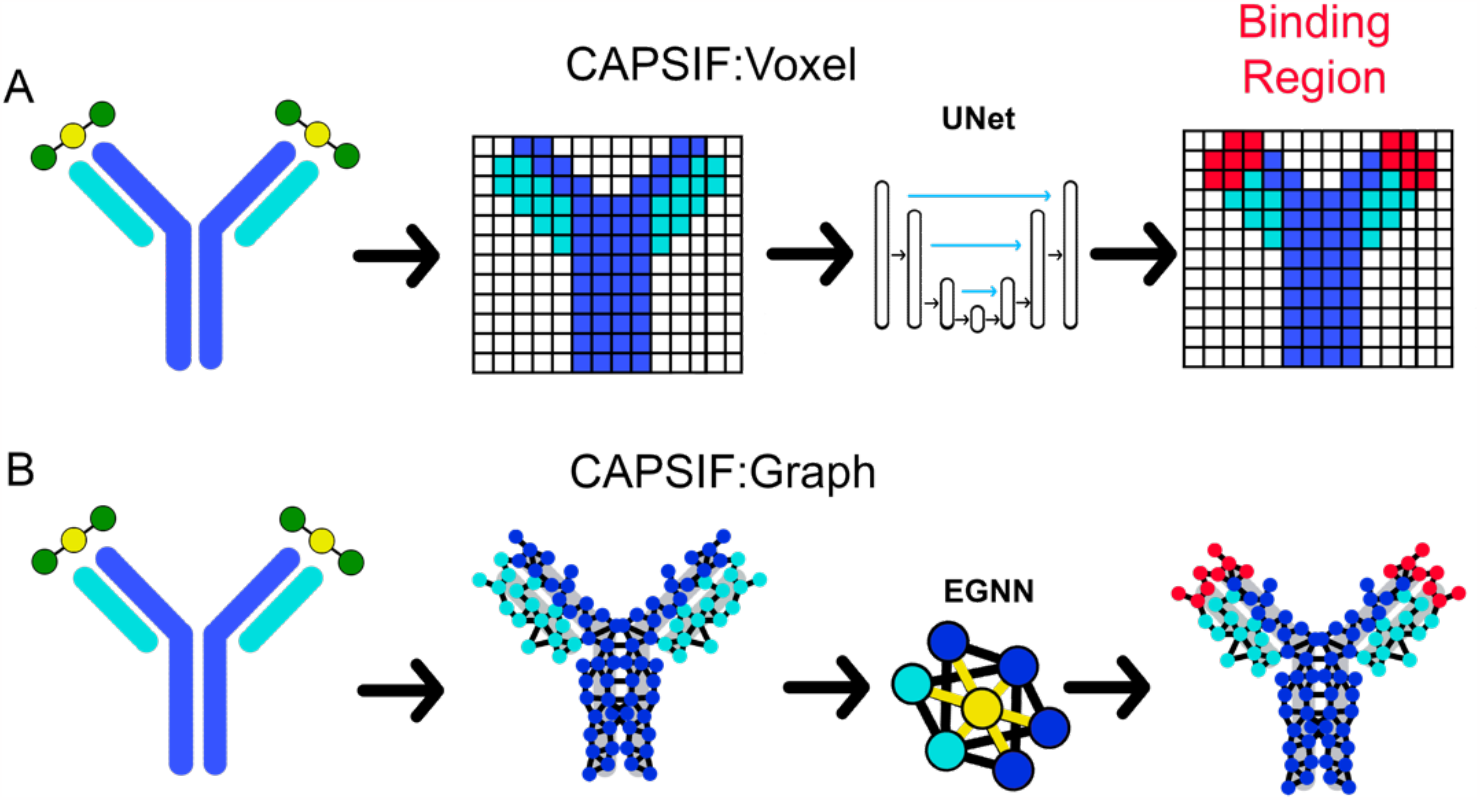
Two deep learning models that predict where proteins bind carbohydrates. **(A)** The first model (CAPSIF:V) maps the β carbon (Cβ) coordinates into voxels, utilizes a convolutional UNet architecture, and predicts the binding residues. **(B)** The second model (CAPSIF:G) converts the Cβ coordinates into network nodes with edges for residue-residue neighbors, performs convolutions on nodes with respect to neighbors with an equivariant graph neural network (EGNN) architecture, and predicts which residues bind sugars.

The second architecture, CAPSIF Graph (CAPSIF:G), is an equivariant graph neural network (EGNN) (50), with each Cβ represented as a node on the graph and edges connected between all neighbor residues within 12 Å (**Figure 1B**). EGNNs use graph-based convolutions with message passing between connected nodes based on node features and the edge features (distances) (50). We explored many variations of these neural network architectures; the Supporting Information includes data supporting our architecture and data representation choices.

The carbohydrate-binding residues comprise 5.2% of the dataset. To ameliorate the effect of data imbalance, we followed Stepniewska-Dziubinska *et al*. and chose the complement of the Dice similarity coefficient (*d*) as our loss function (*L* = 1 − *d*) (21). The Dice coefficient is normalized by both the correctly and incorrectly predicted residues:

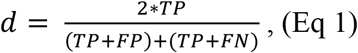

where *TP* = true positives, *FP* = false positives, and *FN* = false negatives. Since *d* does not depend on true negative labels, this loss function is insensitive to imbalanced datasets where the positive label is observed much less than the negative label (21).

### 2.3 CAPSIF predicts carbohydrate-binding residues with encouraging accuracy

CAPSIF:V and CAPSIF:G are novel architectures for predicting carbohydrate binding residues; however, they use 512 structures to train with a substantial data imbalance. We therefore investigated the performance of CAPSIF on a held-out test set to determine whether the architectures accurately predict carbohydrate-binding regions despite the small amount of training data. Four representative CAPSIF:V predictions are shown in **Figure 2**, highlighting *TP* residue predictions, (green), *FP* residues (blue), and *FN* residues (red). CAPSIF:V captures the binding pocket visually for an endoglucanase (**2A**), xylanase (**2B**), and β-glucanase (**2C**), but it performs poorly on the HINT protein that binds ribose (**2D**), a five membered ring carbohydrate that is commonly associated with nucleotides.

**Figure 2:**
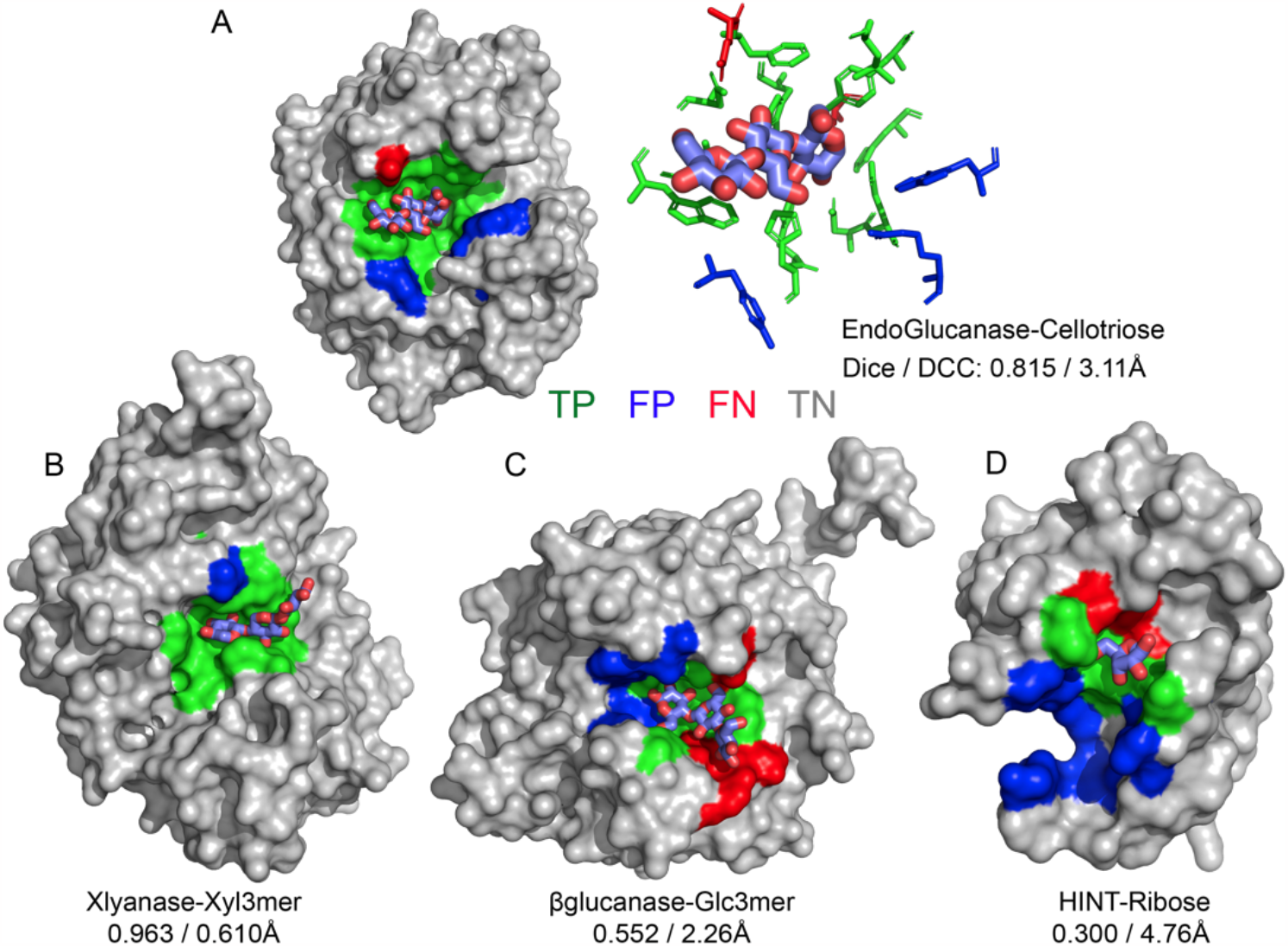
Prediction of carbohydrate binding sites on a protein surface using CAPSIF:Voxel. **(A)** Two representations of binding residues for cellotriose bound to endoglucanase (6GL0), surface (left) and sticks (right); Predicted surface representation of **(B)** xylanase bound to a xylose 3-mer (3W26), **(C) β**-glucanase bound to a glucose 3-mer (5A95), and **(D)** HINT protein bound to a ribose monomer (4RHN) predictions. True positive residue predictions are colored green, false positives are blue, false negatives are red, true negatives are gray, and the bound carbohydrate is cyan; Dice is defined by eq (1) and DCC is distance from center to center of the predicted binding regions.

For comparison, we evaluated how small molecule binding site predictors FTMap (28) and Kalasanty (21) perform for carbohydrate-binding tasks. We assessed these methods using the following metrics: the Dice coefficient (*Eq 1*), distance from the center of the crystal to the center of the predicted binding location (DCC), positive predictive value (PPV), sensitivity, and Matthews correlation coefficient (MCC). Similar to the Dice coefficient, the MCC is suited for unbalanced datasets; it has been reported in previous carbohydrate binding site studies (35–37). MCC is:

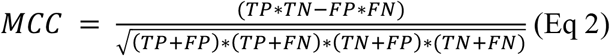

where *TN* = true negatives. MCC ranges from -1 (worst) to +1 (best). The Dice coefficient measures the overlap of correctly predicted interacting residues to all predicted interacting residues. We define a success as a Dice score greater than 0.6 or, following Stepniewska-Dziubinska *et al*., a DCC under 4 Å (21).

On the CAPSIF test set, FTMap achieved an average Dice coefficient of 0.351 and average DCC of 10.5 Å, and Kalasanty achieved an average Dice of 0.108 and average DCC of 14.6 Å (**Table 1**). Further, FTMap predicted 16.8% of test structures with greater than 0.6 Dice and 16.8% of test structures with less than 4 Å DCC, while Kalasanty predicted 0% of test structures with greater than 0.6 Dice and 21.4% of test structures with less than 4 Å DCC (**Table 1, Figure 3A,B**).

**Table 1:** Average metric for each method on test set. Dice similarity coefficient is defined by eq (1), PPV is positive predictive value = TP / (TP + FP), Sensitivity = TP / (TP + FN), DCC is distance from center to center of predicted versus experimentally determined residues and only calculated for proteins that yield predictions (coverage), MCC is Matthews correlation coefficient and defined by eq (2). Bold face indicates best performance for each metric.

| Model | Dice | PPV | Sensitivity | DCC (Å) | MCC | Coverage (%) |
| --- | --- | --- | --- | --- | --- | --- |
| FTMap | 0.351 | 0.284 | 0.505 | 10.56 | 0.222 | **100.0** |
| Kalasanty | 0.108 | 0.080 | 0.207 | 14.62 | -0.624 | 90.0 |
| CAPSIF:V | **0.597** | **0.598** | **0.647** | **4.48** | **0.599** | 94.4 |
| CAPSIF:G | 0.543 | 0.541 | 0.590 | 5.85 | 0.538 | 83.2 |

**Figure 3:**
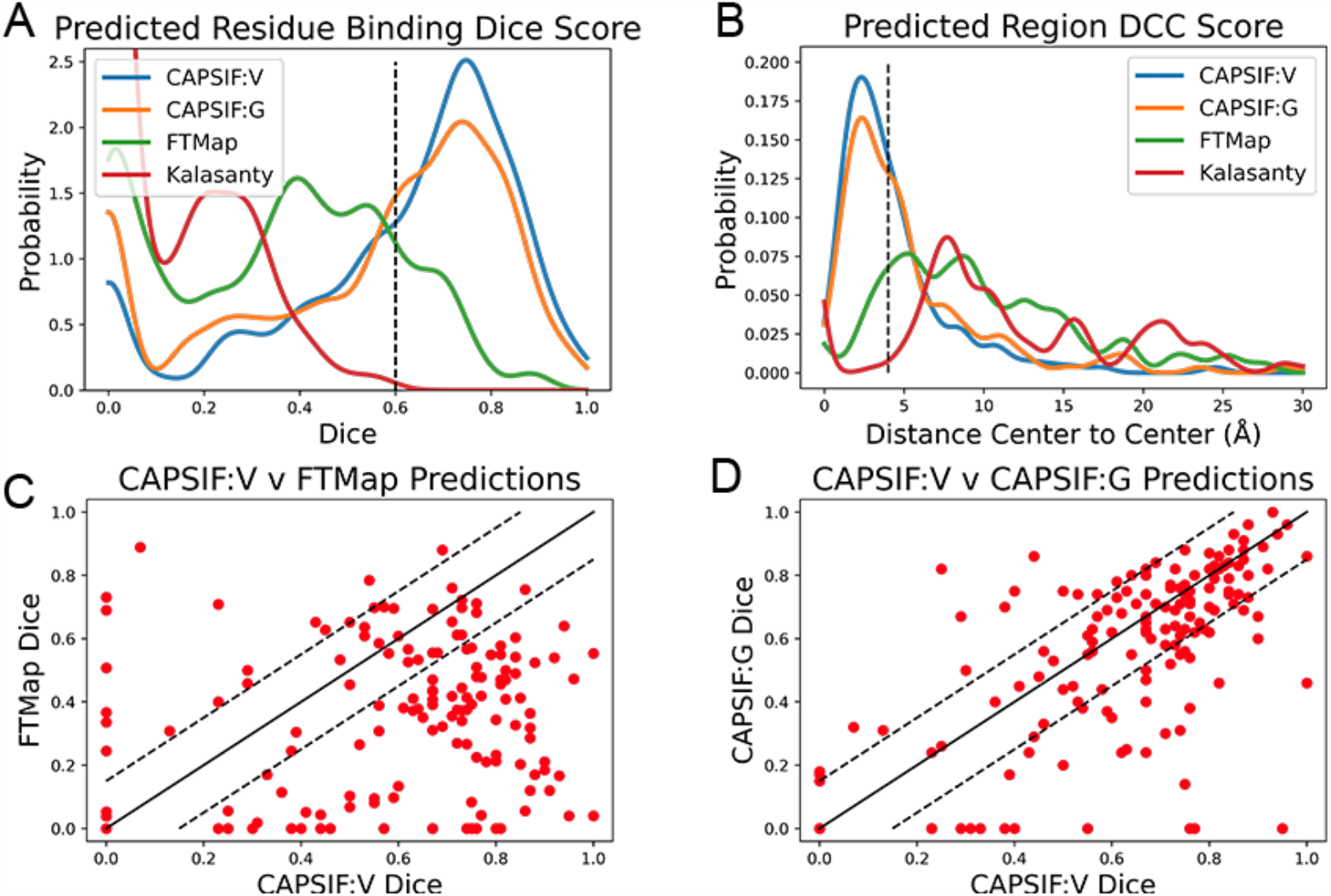
Distributions of CAPSIF:V and CAPSIF:G assessment metrics compared to FTMap (28) and Kalasanty (21). **(A)** Distribution of Dice similarity coefficient for all methods smoothed with a Gaussian kernel density estimate (KDE, bandwidth *h* = 0.04); **(B)** Distance from center to center (DCC) of predicted to experimental carbohydrate binding residues (smoothed with a Gaussian KDE, *h* = 0.75 Å); **(C)** Per-target comparison of CAPSIF:V to FTMap and **(D)** CAPSIF:G Dice coefficients.

We then investigated whether our CAPSIF models, which are specifically tuned for carbohydrate binding, predict the carbohydrate binding regions more accurately than Kalasanty and FTMap. On the held-out CAPSIF test set, CAPSIF:V achieves an average .0596 Dice coefficient and 4.48 Å DCC metric, and CAPSIF:G achieves an average 0.543 Dice coefficient and 5.85 Å DCC metric (**Table 1**). Further CAPSIF:V successfully predicts 62.7% of test structures with greater than 0.6 Dice and 56.5% of test structures with less than 4 Å DCC, and CAPSIF:G successfully predicts 55.2% of test structures with less than 0.6 Dice and 46.0% of test structures with less than 4.0 Å DCC. Both CAPSIF models have a most probable prediction at 0.77 Dice and 2.5 Å DCC (**Table 1, Figure 3A,B**).

Since CAPSIF is ML based and FTMap is energy based, FTMap may predict more accurately on different cases compared to CAPSIF. We compared the CAPSIF:V and FTMap Dice scores for each structure (**Figure 3C**). FTMap achieves a significantly higher Dice coeffiecents (difference greater than 0.15 Dice) than CAPSIF:V in 10.9% of cases, and CAPSIF:V predicts the binding region with a significantly greater Dice coefficient than FTMap in 67.9% of cases. We also compared the computer time. On The FTMap server, FTMap requires an hour or more to predict the binding region for a single structure, whereas both CAPSIF:V and CAPSIF:G predict binding sites within seconds on a single CPU. Thus, on average, CAPSIF:V and CAPSIF:G outperform current small molecule binding site predictors for carbohydrate binding.

Finally, we compared the CAPSIF:V architecture to the CAPSIF:G architecture. CAPSIF:V has an average Dice coefficient of 0.596 and CAPSIF:G has an average Dice coefficient of 0.543 across the test dataset (**Table 1**). When comparing the Dice on the test set, CAPSIF:V predicts 27.3% of structures with greater than 0.15 Dice than CAPSIF:G, while CAPSIF:G predicts 11.2% of structures with greater than 0.15 Dice than CAPSIF:V (**Figure 3D**). Thus, CAPSIF:V outperforms CAPSIF:G on carbohydrate binding site prediction.

Carbohydrates are unique biomolecules that bind to different lectins with high specificity. Both CAPSIF architectures treat all carbohydrates agnostically, meaning that all sugar residue types are considered equivalent for predictions. Nonetheless, we compared prediction results across different sugar residue types. (**File SI1**). CAPSIF:V performs best on glucose (Glc), galactosamine (GalN), arabinose (Ara), xylose (Xyl), ribose (Rib), and galacturonic acid (GalNAc). It predicts regions that bind neuraminic acid (Neu/Sia), fucose (Fuc), and Glucuronic acid (GlcNAc) with less than an average 0.5 Dice coefficient. The weaker performance could stem from the chemical differences or differences in the size of the training data. Neu and Fuc are substantially chemically distinct carbohydrates, as Neu is a 9-carbon structure and Fuc adopts an (*L*) conformation; both are sparse in our dataset.

### 2.4 CAPSIF:Voxel performs equivalently on AlphaFold2 structures

Both CAPSIF models were trained and tested on bound crystal structures; however, experimental protein structure determination can be expensive, even in the absence of a carbohydrate. We therefore investigated whether CAPSIF:V could usefully predict carbohydrate binding structures from computationally modeled structures. We reconstructed the test protein structure dataset with the Colab implementation of AlphaFold2 (AF2) (20, 51) and predicted the carbohydrate binding residues of the predicted structures and evaluated the same performance metrics (**Table 2**). CAPSIF:V predicts the carbohydrate binding regions with similar Dice coefficients of 0.597 and 0.586 for protein databank versus AF2 predicted structures, respectively. **Figure 4A** shows that the Dice distribution is similar between PDB and AF2 structures. CAPSIF:V predicts the center of the carbohydrate binding region more accurately on AF2 structures with a DCC of 3.8 Å, compared to 4.5 Å on crystal structures.

**Table 2:** Metrics for CAPSIF:Voxel inputting PDB or AF2 structures. Dice, PPV, Sensitivity, DCC, MCC, and defined in Table 1.

| Structures | Dice | PPV | Sensitivity | DCC (Å) | MCC | Coverage (%) |
| --- | --- | --- | --- | --- | --- | --- |
| PDB | 0.597 | 0.598 | 0.647 | 4.48 | 0.599 | 94.4 |
| AF2 | 0.586 | 0.508 | 0.744 | 3.76 | 0.598 | 85.0 |

**Figure 4:**
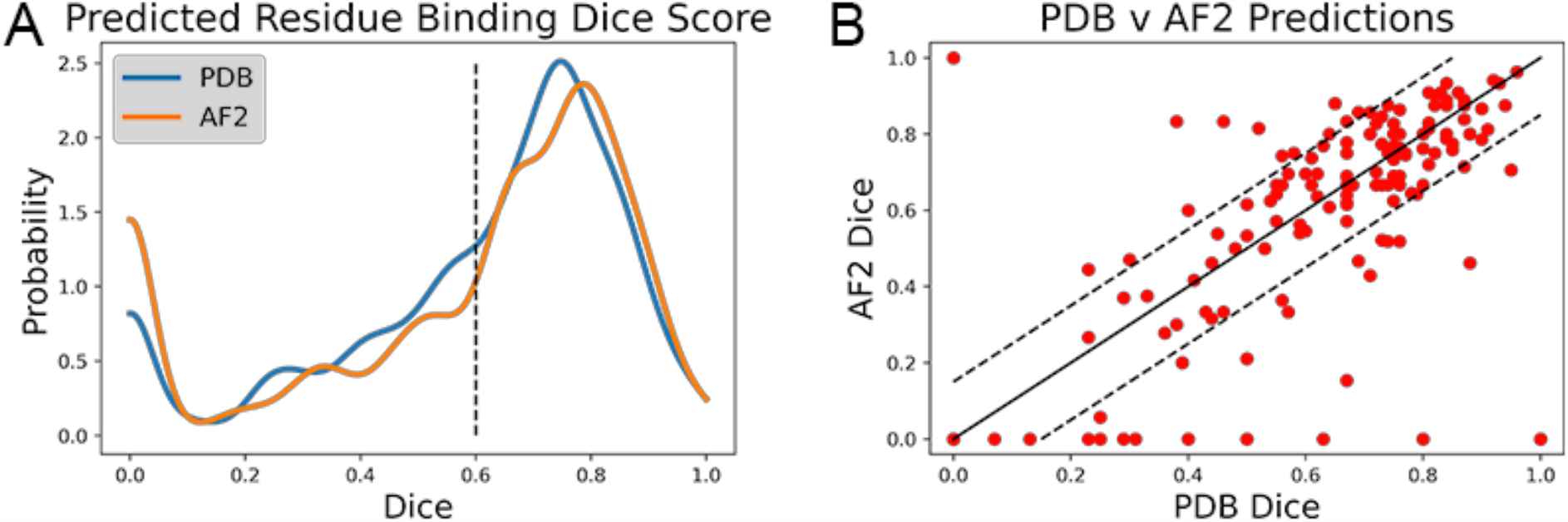
Dice coefficient assessment of CAPSIF:Voxel on PDB and AlphaFold 2 (AF2) structures. **(A)** Kernel density estimate (*h* = 0.04) showing the distribution of Dice coefficient for both methods; **(B)** Comparison of each test structure between CAPSIF:V on PDB and AF2 structures.

Although CAPSIF:V has a lower average DCC on AF2 structures compared to experimental structures, CAPSIF:V fails to predict any sites at all on 15% of AF2 structures, whereas it fails in only 5% of PDB structures.

The multiple outliers where CAPSIF:V fails to predict the region of carbohydrate binding in only AF2 predicted structures are sorted in **Figure 4B**. CAPSIF:V predicts a Dice coefficient of at least 0.15 units higher for PDB structures in 14.9% of structures and predicts AF2 structures with a 0.15 Dice coefficient or higher for 8.7% of test structures. AF2 generated structures can be inaccurate; however, in most of the test cases, AF2 captures the structures with angstrom level accuracy and the carbohydrate binding residues with high pLDDT confidence; unfortunately the pLDDT confidence measure does not correlate with the CAPSIF success rate (**Figure S8**).

### 2.5 CAPSIF assists *ab initio* prediction of bound protein-carbohydrate structures

CAPSIF:V predicts the carbohydrate binding site on the majority of proteins with high accuracy, suggesting that it might be used in a pipeline to predict bound protein-carbohydrate structures. As a proof-of-concept, we developed a prospective pipeline and tested it on five proteins from the GlycanDock (24) test dataset that were not included the CAPSIF dataset.

We constructed the following rudimentary pipeline. We predicted the binding site from each unbound protein’s experimentally determined structure with CAPSIF:V and constructed the known carbohydrate with Rosetta. The carbohydrate’s center of mass (CoM) was then placed in the CoM of the predicted binding region and manually rotated to align with the binding region shape. Next, we used the Rosetta FastRelax (52) protocol to remove steric clashes. Then we used Rosetta’s standard GlycanDock (24) to predict the bound structures. To find the highest rated bound structure, we filtered 9,500 decoys by their computed interaction energy.

We tested the pipeline on five experimentally solved unbound proteins: *P. aeruginosa* lectin 1, a glycan binding protein (GBP, 1L7L), two carbohydrate binding modules (CBMs, 1GMM and 2ZEW), a glycoside hydrolase enzyme (1OLR), and an anti-HIV-1 antibody (Ab) (6N32). **Figure 5** shows structures and the root mean squared deviation (RMSD) of each predicted carbohydrate structure from the experimental structure. CAPSIF:V predicted carbohydrate binding residues near the correct site on four of the five proteins, but it failed to predict any binding residues on the antibody (6N32). For three of the proteins, CAPSIF:V predicts the region with high accuracy, but on 1GMM, CAPSIF:V predicts regions flanking the binding site, but still provides a similar CoM to the actual binding region. For the for carbohydrates with identified sites, the standard GlycanDock protocol was able to refine the carbohydrate structure to an RMSD of less than 8 Å for the entire ligand and less than 6 Å for register-adjusted values, where the termini were removed before calculating RMSD. The 3-mer Gal GBP (1L7L) has the worst RMSD (6 Å register adjusted, **Figure 5B**), likely because the *holo* conformation (2VXJ) undergoes a conformational change at the carbohydrate-binding site. These predictions demonstrate the potential of CAPSIF to help inform experimental hypotheses or for high throughput predictions of bound protein-carbohydrate structures.

**Figure 5:**
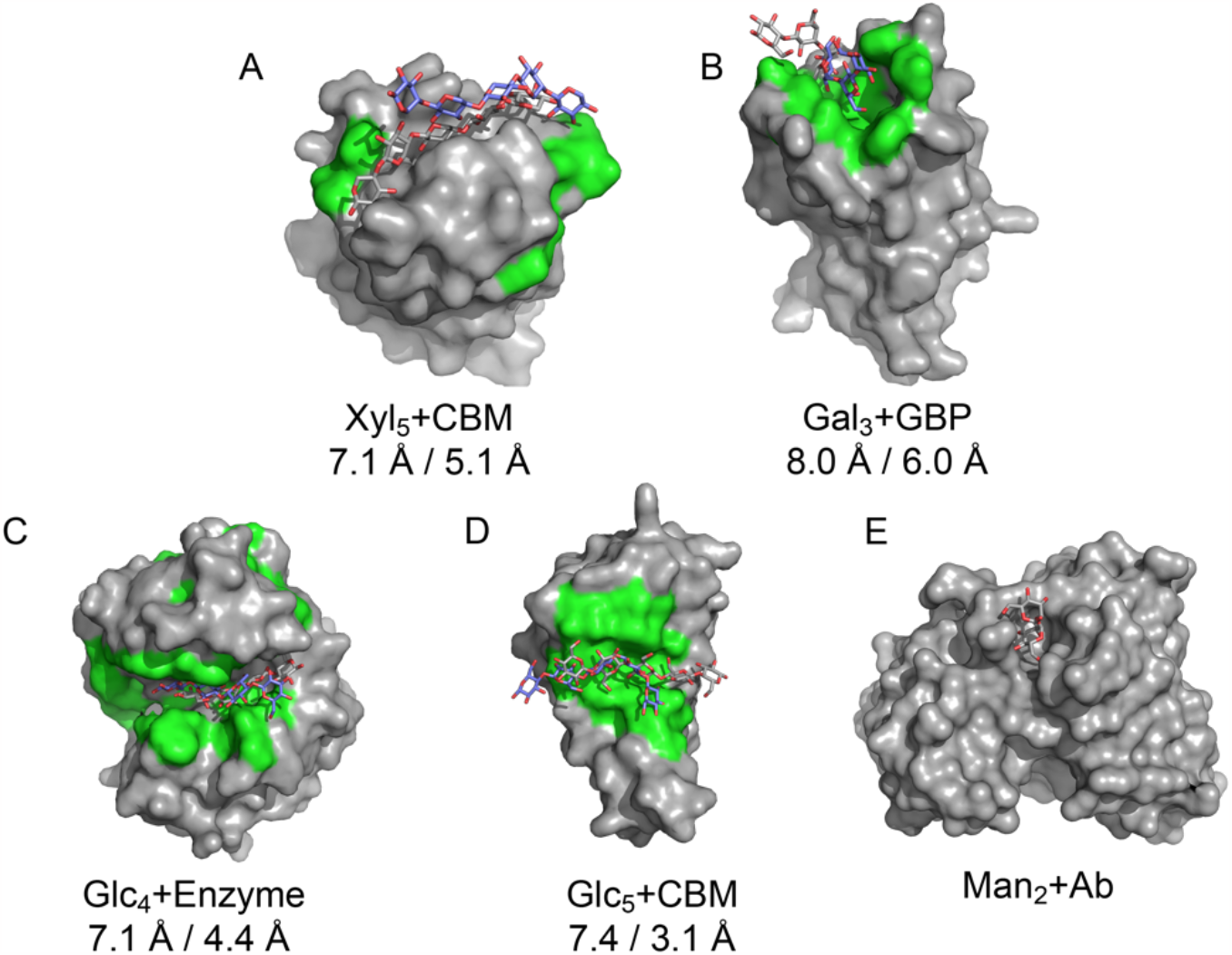
Results of CAPSIF:V-GlycanDock pipeline. CAPSIF-predicted residues are shown in green. Wild type unbound structures are shown in surface representation in gray with the experimentally determined carbohydrate in gray sticks and predicted bound carbohydrate in purple sticks. RMSD of entire ligand and RMSD of register-adjusted ligand are shown below. **(A)** a carbohydrate binding module (CBM), 1GMM (unbound PDB)/1UXX (bound PDB), **(B)** a glycan binding protein (GBP), 1L7L/2VXJ, **(C)** an enzyme, 1OLR/1UU6, **(D)** a CBM, 2ZEW/2ZEX, and **(E) an** antibody (Ab), 6N32/6N35.

## 3 Discussion

We demonstrated that both CAPSIF models predict residues of proteins that bind carbohydrates with much higher accuracy than prior approaches. CAPSIF:V uses a voxelized approach and predicts 62.7% of crystal structures with a distance from the center of the predicted region to the center of the experimentally determined region (DCC) within 4 Å. CAPSIF:G performs strongly on the dataset predicting 55.2% of crystal structures with a DCC less than 4 Å, with CAPSIF:V performing similarly or outperforming CAPSIF:G in most cases. CAPSIF:V is robust to errors in protein structure of the magnitude in AF2 structures: the algorithm predicts similar carbohydrate-binding residue regions independent of whether the input structure is experimentally determined or predicted by AF2. This algorithm is a substantial improvement over surrogate ligand site predictors Kalasanty and FTMap.

Further, CAPSIF outperforms previous methods specifically tuned for carbohydrate binding. CAPSIF:V achieves a 0.599 MCC and CAPSIF:G achieved a 0.538 MCC on the test dataset. Tsia *et al*’s method using probability density maps achieved a 0.45 MCC on their independent test dataset of 108 proteins (35), SPOT-Struc achieved a ∼0.45 MCC on their test dataset of 14 proteins (36), and SPRINT-CBH achieves a MCC of 0.27 MCC on their test set of 158 proteins (37). While these datasets differ from ours, ours is a similarly constructed non-homologous dataset of 162 structures, and CAPSIF has markedly stronger MCC.

Although CAPSIF accurately captures the protein-carbohydrate binding interface, there are limitations. CAPSIF is carbohydrate-agnostic, so it only predicts that a protein residue will bind one of 16 carbohydrate monomers. That is, CAPSIF predicts the location of carbohydrate binding but not which carbohydrate preferentially binds there. Further, CAPSIF was only trained and tested on known carbohydrate binding proteins, therefore CAPSIF may not be informative on non-carbohydrate binding proteins. Another limitation is that CAPSIF fails to predict any binding on about three times as many AF2 predicted structures as crystal structures. Unfortunately, CAPSIF prediction accuracy on AF2 structures is not correlated with pLDDT confidence metrics so it is not possible to know when it will fail.

The scope of CAPSIF makes it well suited for a computational pipeline. We suggest the use of DeepFRI (53), a deep learning model that predicts protein function, to first determine if the protein is a carbohydrate binding protein. If the protein is a carbohydrate binding protein, then LectinOracle(41) or GlyNet (42) can be used to predict which carbohydrates bind the protein. CAPSIF can then predict binding locations, either from an experimental structure or AF2 generated structures, and then GlycanDock(24) can predict a docked protein-carbohydrate structure.

We tested part of this pipeline by predicting the binding region using CAPSIF:V and docking the known carbohydrate binder to the region with GlycanDock (24). CAPSIF:V predicted binding sites on four of the five proteins. The antibody case, which failed, binds a carbohydrate at the complementary determining region (CDR) loops, split over two chains, but CAPSIF was trained only on single chain data. When register adjusted, each structure yielded a ligand RMSD less than 6 Å. We anticipate that a more well-tuned pipeline could yield higher accuracy structures *ab initio* from sequence only.

To our knowledge, voxelized and graph-based site prediction has not been presented simultaneously before. Existing studies have used graphs to either predict binding affinity (54) or a docked structure (in coordination with diffusion techniques) (55), but they have not been used to determine small molecule binding regions. We tested two architectures utilizing either voxel or graph representations. We showed that CAPSIF:V outperforms CAPSIF:G, both of which use convolutions to predict the carbohydrate binding ability of residues with the same residue representation. We can speculate about the reason by considering the differences between the approaches. CAPSIF:V discretizes the protein structure over a 3D grid, which can obscure the Cβ position by a few Ångströms, whereas CAPSIF:G uses the coordinates without any loss of spatial information. CAPSIF:V encodes the initial ∼1.4M feature input to a lower dimensionality of a 512-feature vector to encode the entire structure, whereas CAPSIF:G lifts the data from an *N*_res_ x 30 to a higher dimensionality of *N*_res_ x 64. CAPSIF:V has ∼102M parameters and CAPSIF:G has ∼236K parameters, reflecting how graph-based methods capture the spatially equivariant information in fewer parameters. One characteristic of using the voxel representation is that the grid contains voxels with the protein and the voxels outside the protein, including binding pocket cavities, whereas the graph representation only contains the protein. The voxel network reasoning over the surface pocket volume may be the key factor for improved carbohydrate-binding residue prediction.

Building on this initial screen, future studies could focus on improving the CAPSIF data representation for improved accuracy and extending these models to predict which carbohydrate monomer a residue most preferentially binds as well as whether the protein is a carbohydrate-binding protein. Although lectins are well known for carbohydrate binding, some protein families, such as G protein coupled receptors (GPCRs) and antibodies, do not exclusively bind carbohydrates (56, 57). High throughput methods like these could enable proteomic scale sorting of carbohydrate binding capabilities.

## 4 Methods

### 4.1 Dataset

No dataset of nonhomologous bound protein-carbohydrate structures existed that leveraged the total size of the current PDB, so we constructed one. Simply selecting all RCSB (43) structures with carbohydrates gives all docked protein-carbohydrate structures but also inherently returns all glycosylated proteins, glycosylated peptides, as well as all protein structures that use carbohydrates as crystallization agents. We desired to determine all true physiological protein-carbohydrate interactions, so therefore we manually removed nonspecific crystallization buffers or glycoproteins. Next, we removed all proteins with resolution over 3 Å. Then we removed all homologous protein structures over 30% sequence identity to remove all sequentially redundant proteins. Some structures containing sugars with modified monosaccharides and cyclic carbohydrates were unreadable in the PyRosetta (58) software and therefore additionally removed.

The final dataset consists of 808 structures, with a split of 521 training structures, 125 validation structures, and 162 test structures. Each structure has one or more of the following carbohydrate monomers: glucose (Glc), glucosamine (GlcNAc), glucuronic acid (GlcA), fucose (Fuc), mannose (Man), mannosamine (ManNAc), galactose (Gal), galactosamine (GalNAc), galacturonic acid (GalA), neuraminic acid (Neu)/sialic acid (Sia), arabinose (Ara), xylose (Xyl), ribose, rhamnose (Rha), abequose (Abe), and fructose (Fru). The numbers of each monomer per structure and Dice coefficient for each carbohydrate monomer type from CAPSIF:V are included in **Supplementary File S1**. For all following work, we defined a carbohydrate-interacting residue as residues with any heavy atom that is within 4.2 Å of a carbohydrate heavy atom.

### 4.2 CAPSIF:V Data Processing

Convolutional neural networks are not rotation invariant, and so data augmentation by rotations improves their performance (59). Therefore, we augmented the input data for CAPSIF:V during training to overcome the rotational variance. Each time a structure was used in training, it was rotated in Cartesian space by a random angle in {-180°,180°} around an axis defined by a randomly-chosen residue’s location and the protein center-of-mass. With the random rotation for each epoch, the network learned approximately 1,000 different orientations of each structure in the data set. If the protein was too large for the grid size, the protein was split into separate grids and run separately (about 22% of the training points).

### 4.3 Neural Network Architectures

#### 4.3.1 Features

Due to the small dataset size of 808 structures, we chose residue-level representations instead of atomistic. We assigned all residue information to the Cβ atom of each residue because the position of the Cβ is similar in *apo* and *holo* states (47). The features are listed in **Table 3**. The SASA, hydrophobicity, H bond donor/acceptor indices were calculated using pyRosetta (58), and aromatophilicty was indexed by Hirano and Kameda (49).

**Table 3:** List of features and the associated encoding size used for both CAPSIF models.

| Feature Type | Encoding Size |
| --- | --- |
| Amino acid (one-hot) | 20 |
| SASA | 1 |
| Hydrophobicity | 1 |
| Aromatophilicity | 1 |
| H Bond Donor/Acceptor | 2 |
| Orientation (Voxel only) | 3 |
| Torsion (Graph only) | 4 |

#### 4.3.2 CAPSIF:Voxel

CAPSIF:V utilizes a UNet architecture, encoding and decoding the input structure to predict carbohydrate binding residues with residual connections. CAPSIF:V inputs a grid of 36 × 36 × 36 voxels with each voxel representing 2 Å × 2 Å × 2 Å. We input a tensor of size (28,36,36,36), with the 28 features from **Table 3**, where orientation is the normalized components of the Cα to Cβ bond vector. All voxels without a Cβ within are input as zero-vectors.

CAPSIF:V contains an embedding layer and 9 convolutional blocks where 4 blocks encode the structure, 1 block forms the bottleneck, and 4 blocks decode the structural information. The embedding layer lifts the 28-channel input into a 32-dimension space. Each block has a double convolution, performing the following methods twice: 3D convolution, with the same number of input channels as number of output channels, (5×5×5) kernel with a stride of 1 and padding of 2, a batch normalization layer, and rectified linear units (ReLU) activation function. In addition, each encoding block also has a MaxPooling layer to double the size of the channels (32,64,128,256,512) while reducing 3D cubic voxel number (36,18,9,3,1). Each decoding block first concatenates the results of the encoding layer of the same size and then performs a double convolution and a 3D-transposed convolution operator, reducing the number of channels (256,128,64,32) while increasing the 3D cubic voxel number (3,9,18,36). After the 9 blocks, there is a single convolutional layer condensing the input channels (32) into a single output channel, which is then followed by a sigmoid activation function to output the probability that the voxel contains a residue that binds a sugar (**Figure 6**). CAPSIF:V contains 102,676,001 parameters.

**Figure 6:**
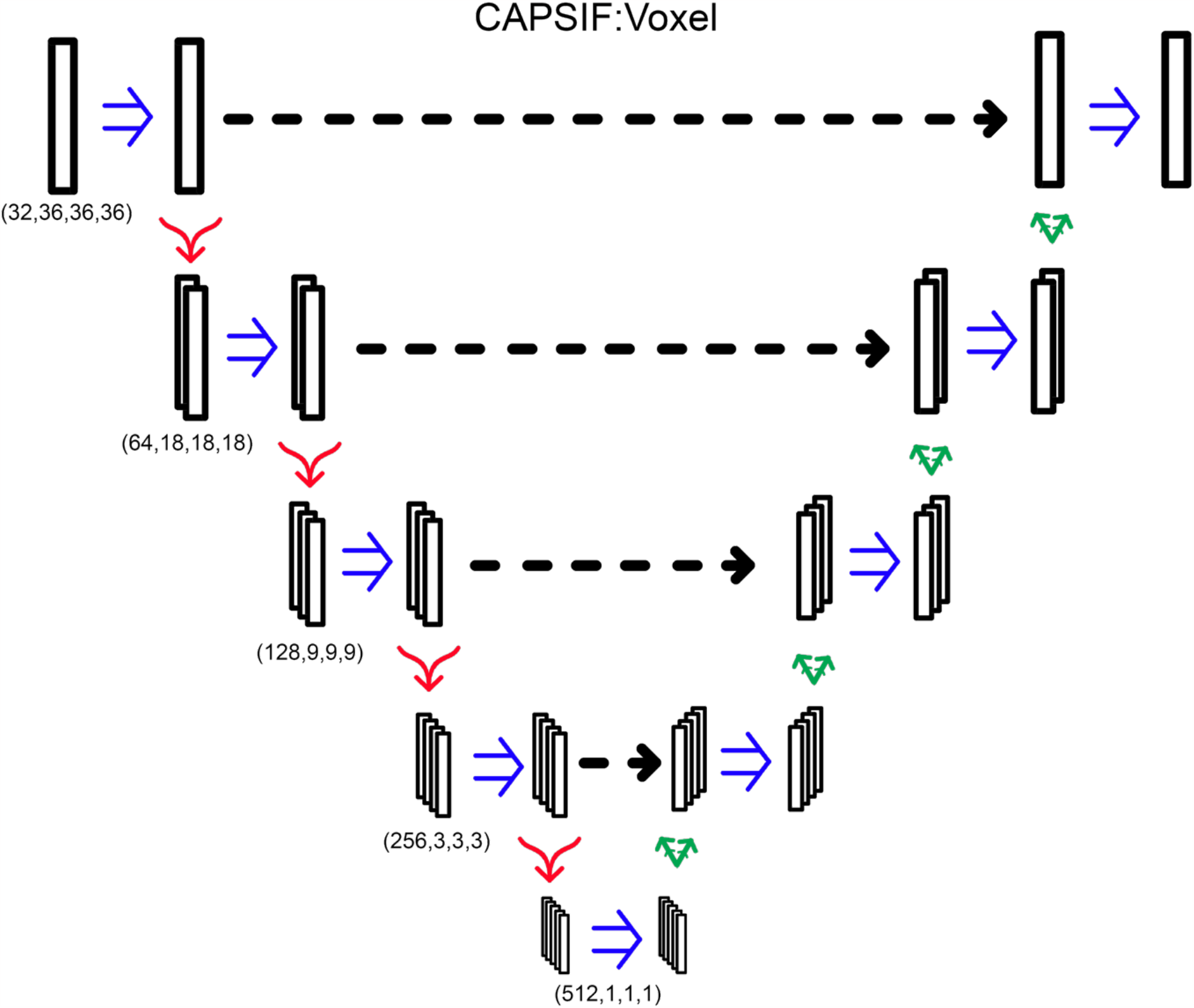
CAPSIF:V architecture. Blue arrows indicate a double convolution, red arrows indicate an encoding layer, and green arrows indicate a decoding layer.

CAPSIF:V was trained for 1,000 epochs with a learning rate of 10^−4^ and batch size of 20 grids using the Adam (60) optimizer with the loss function *L* = 1 − *d*, where *d* is defined by (*Eq 1*).

#### 4.3.3 CAPSIF:EGNN

CAPSIF:G is an equivariant graph neural network (50) that performs convolutions on each node (chosen as each Cα for glycine and Cβ for all others). Graph edges are connected between neighbors (defined as all other nodes’ within 12 Å) and the edge attribute is the distance between node Cβ atoms. In addition to the features used in CAPSIF:V, we include a torsional component in the node features as the sine and cosine of the φ and ψ angles of each residue (**Table 3**).

CAPSIF:G first lifts the 29-feature input node into a 64-dimension space. The 64-feature vector, alongside the edge features (distances) is then input to eight consecutive equivariant graph convolutional layers (EGCLs) (50). Each EGCL contains an edge multilayer perceptron (MLP), a node MLP, a coordinate MLP, and attention MLP. The edge MLP consists of two blocks of a linear layer and a rectified linear units (ReLU) activation function. The node MLP consists of a linear layer, a ReLU activation layer, and linear layer. The coordinate MLP contains a linear layer, a ReLU activation layer, and a linear layer. The attention MLP contains a linear layer and a sigmoid activation function. All layers input and output a 64-feature vector. Finally, CAPSIF returns the embedding to a 29-feature vector per node, adds the initial input features to the final vector, performs batch normalization, and then uses a sigmoid activation function to output a probability of carbohydrate binding of all residues. CAPSIF:G contains 236,009 parameters.

This model was trained for 1,000 epochs with a learning rate of 10^−4^ and batch size of one protein using the Adam optimizer (60) with the loss function *L* = 1 - *d*, where *d* is defined by (*Eq 1*).

## 5 Data Availability Statement

The datasets and the code for each model are available for non-commercial use at https://github.com/Graylab/CAPSIF.

## 6 Author Contributions

S.W.C. wrote the text and created figures, explored variations of the CAPSIF:EGNN model, and analyzed data. S.S. conceptualized the project, created the models and the dataset, analyzed data, and wrote an initial manuscript. J.J.G. conceived and supervised the project, analyzed data, and wrote the text.

## 8 Acknowledgements

S.W.C. and S.S. thank members of the Gray lab for insightful discussions, notably Sai Pooja Mahajan, Rituparna Samanta, and Jeffrey Ruffolo. Computational resources were provided by the Advanced Research Computing at Hopkins (ARCH).

