## Supplementary Information for "Structure-Based Neural Network Protein-Carbohydrate Interaction Predictions at the Residue Level"

### Dataset

The dataset is composed of the following monomers: glucose (Glc), glucosamine (GlcNAc), glucuronic acid (GlcA), fucose (Fuc), mannose (Man), mannosamine (ManNAc), galactose (Gal), galactosamine (GalNAc), galacturonic acid (GalA), neuraminic acid/sialic acid (Neu/Sia), arabinose (Ara), xylose (Xyl), ribose, rhamnose (Rha), abequose (Abe), and fructose (Fru). There are either no training structures or very few for Fru, ManNAc, Abe, Rha, ribose, GalA, and GlcA. Although some have a high average test Dice similarity coefficient, CAPSIF may not accurately predict protein residues that bind those carbohydrate species well. Finally, CAPSIF:Voxel does not perform well on predicting residues that bind Neu and Fuc, likely due to their 9-carbon structure and (*L*) conformation, respectively, as well as GlcNAc.

The supporting Excel file **Supplementary File S1** includes the following information:

- PDB ID
- Carbohydrate species
- Per-PDB CAPSIF:V Dice coefficient
- Per-carbohydrate species Dice coefficient

### Determination of Data Representation

For voxel locations, we compared three representation choices, (1) α carbon (Cα), (2) β carbon (Cβ), or (3) Cα and Cβ positions for the location of voxels. We trained and tested each of these models as described in the Methods. We compared the Dice coefficient, sensitivity and positive predictive value to determine which representation performs best (**Figure S1**, **Table S1**). The Cβ-only representation has an average test Dice coefficient of 0.551, with the Cα representation having a test Dice coefficient of 0.545, where when both the Cα and Cβ are included together in the representation, the architecture has an average test Dice coefficient of only 0.528.

Finally, we further included orientation information of the residues themselves by concatenating the unit vector of the Cα to Cβ bond to the Cβ only representation. This representation had an average test metric of 0.597 (Cβ: Cα → Cβ vec) (**Figure S1**, **Table S1**). This method performed the best of all three representations, having the largest coverage and highest average test metrics. For these reasons, we chose Cβ: Cα → Cβ as our representation of coordinates and orientation for CAPSIF:V.

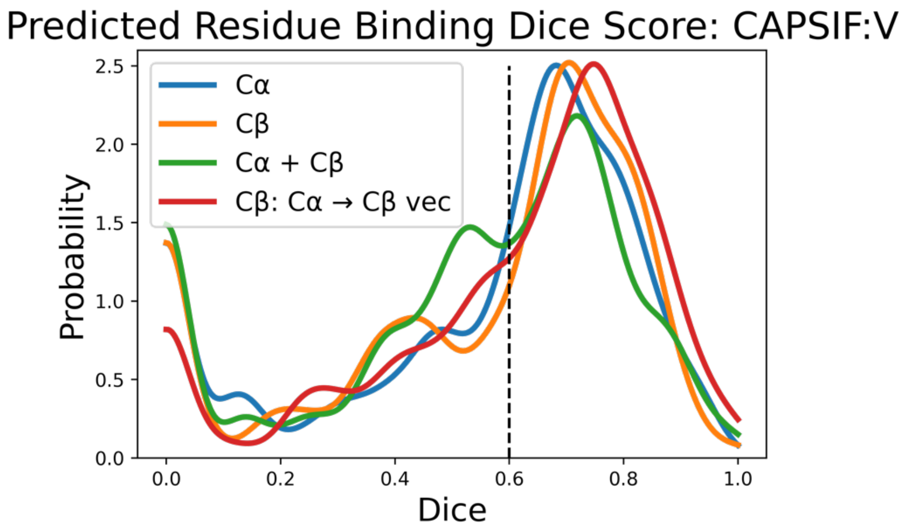


**Figure S1**: **Test Dice coefficient assessment for different representations with CAPSIF:V architectures**: Blue shows a Cβ representation including a normalized vector for alpha carbon (Cα) to Cβ, orange shows only a Cβ representation, green shows Cα representation, and red shows Cα and Cβ representation with all voxels.

**Table S1**: **Performance for each CAPSIF:V model**. Dice coefficient is defined by (*Eq 1)*; PPV and Sensitivity are same as Table 1.

| **Voxel Representation** | **Dice** | **PPV** | **Sensitivity** |
|---|---|---|---|
| Cβ | 0.551 | 0.563 | 0.583 |
| Cα | 0.545 | 0.535 | 0.620 |
| Cα + Cβ | 0.528 | 0.555 | 0.554 |
| Cβ: Cα → Cβ | **0.597** | **0.598** | **0.647** |

Next, we investigated CAPSIF:G node representations, with the architecture described in Methods. We constructed the following variants: Cβ nodes with φ and ψ angles, Cβ and N, Cα, and C backbone nodes (and one-hot encoding for atom type, without φ and ψ angles). The Cβ only node representation performed the best with a Dice coefficient of 0.543. Further, Cβ takes a fraction of the time for predictions compared to the backbone due to graph construction time, therefore we chose the CAPSIF:G to be the Cβ model (**Figure SI2, Table SI2**).

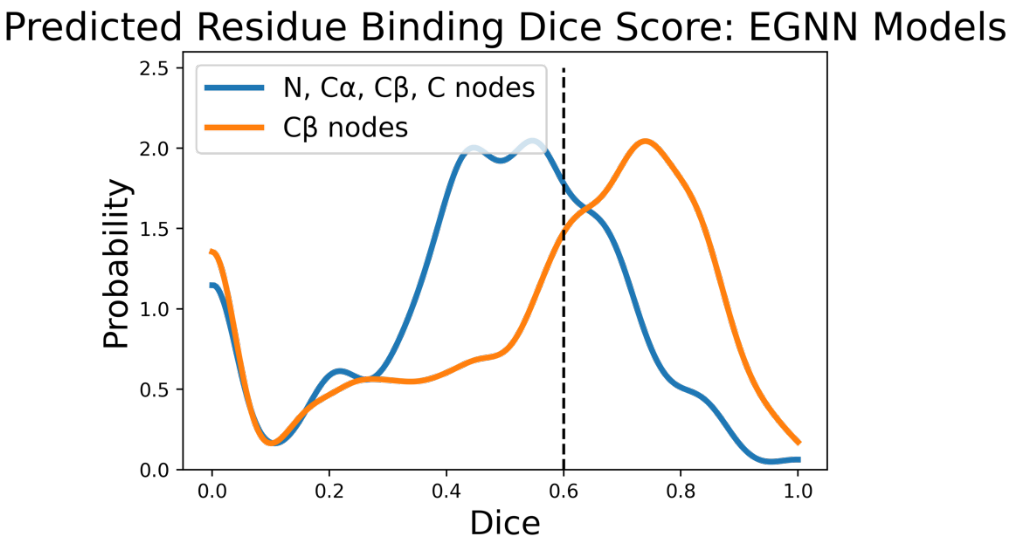


**Figure S2: Test Dice coefficient assessment for different representations with EGNN architectures**. Blue shows all backbone atoms node representation, orange shows a Cβ node representation.

**Table S2**: **Performance for EGNN model node representation**. Dice coefficient is defined by (*Eq 1)*; PPV and Sensitivity are same as Table 1.

| **EGNN Representation** | **Dice** | **PPV** | **Sensitivity** |
|---|---|---|---|
| Cβ | **0.543** | **0.541** | **0.590** |
| Backbone | 0.458 | 0.396 | 0.647 |

### Random Assignment of Carbohydrate Binding Regions

As a control, we compared CAPSIF to a random baseline. For example, for 200 amino acids with a 5.0% positivity rate, we randomly select 10 residues as a true label (sugar binding) and computed the Dice similarity coefficient (*Eq 1*). Using 1,000 trials for an endoglucanase (6GL0), which has 331 total residues with 14 that experimentally bind carbohydrates, we observe a theoretical maximum Dice coefficient at approximately 0.08 when all residues are predicted as carbohydrate binders. At a rate of 5%, we observe a mean Dice coefficient of 0.046, where CAPSIF:V predicts that protein with a Dice coefficient of 0.963 (**Fig SI 3A**).

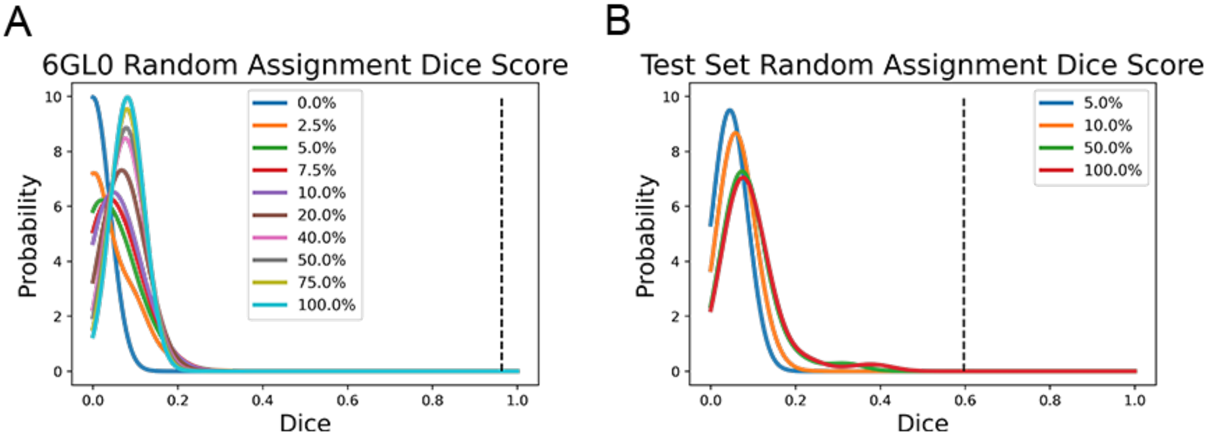


**Figure S3**: **Dice coefficient assessment with random assignment smoothed with a kernel density estimate with bandwidth *h* = .04**. **(A)** Dice evaluation of random assignment of an endoglucanase (6GL0). **(B)** Dice evaluation over entire test set.

The dataset has, on average, 5.16% of protein residues bind carbohydrates. With random assignment over the entire dataset, random assignment at 5.16% yields an average 0.046 Dice score, where CAPSIF:V outperforms random assignment by over 12-fold at an average 0.593 Dice (**Fig SI4B**).

### Determination of CAPSIF probability threshold

To determine the best probability cutoff value for the final activation function, we altered the threshold on the test dataset (**Fig SI5**). CAPSIF:V differs minimally for all thresholds while CAPSIF:G negatively correlates with increasing threshold and drops more sharply after a cutoff of 0.6. For both architectures we chose a threshold of 0.5.

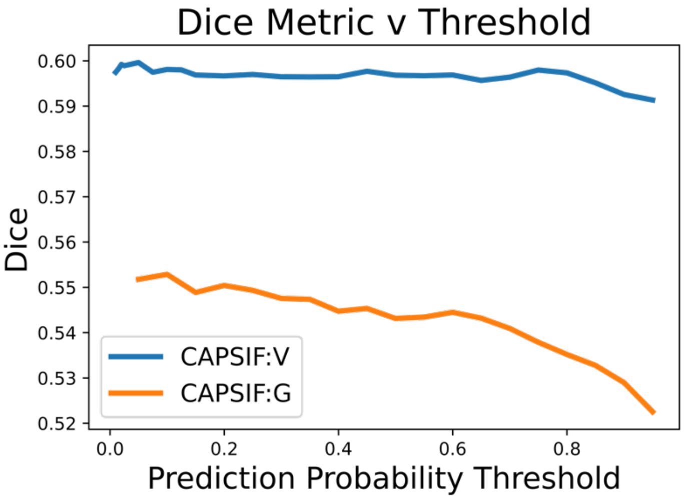


**Figure S4: Test Dice coefficient assessment for CAPSIF architectures for various thresholds for the final sigmoid activation function**. Blue represents CAPSIF:V, orange represents CAPSIF:G.

### Comparison of Dice and DCC metrics

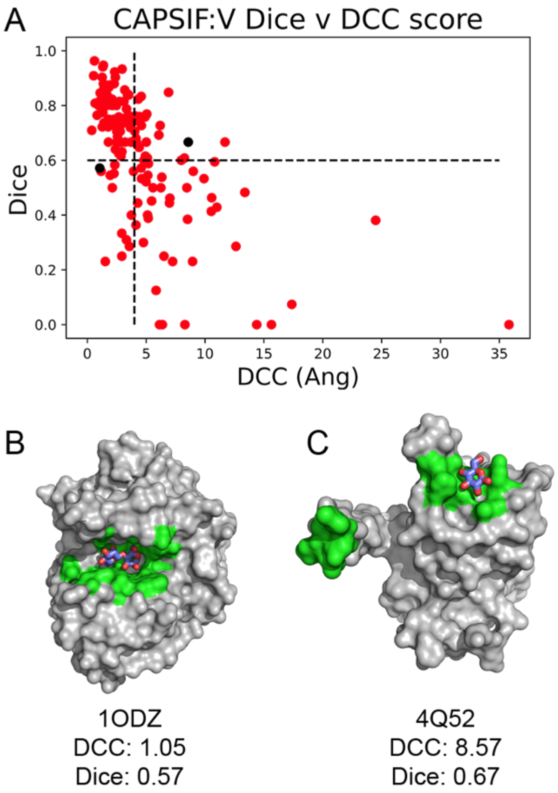


**Figure S5**: **Comparison of Dice score and DCC**. **(A)** Per-target comparison of Dice and DCC for CAPSIF:V predictions on the test set. CAPSIF:V predictions (green) on **(B)** endo-1,4-β-mannosidase 1ODZ and **(C)** *C. pinesis* DSM 2588 (4Q52) (gray).

### Figures comparing CAPSIF:Voxel and CAPSIF:Graph predictions

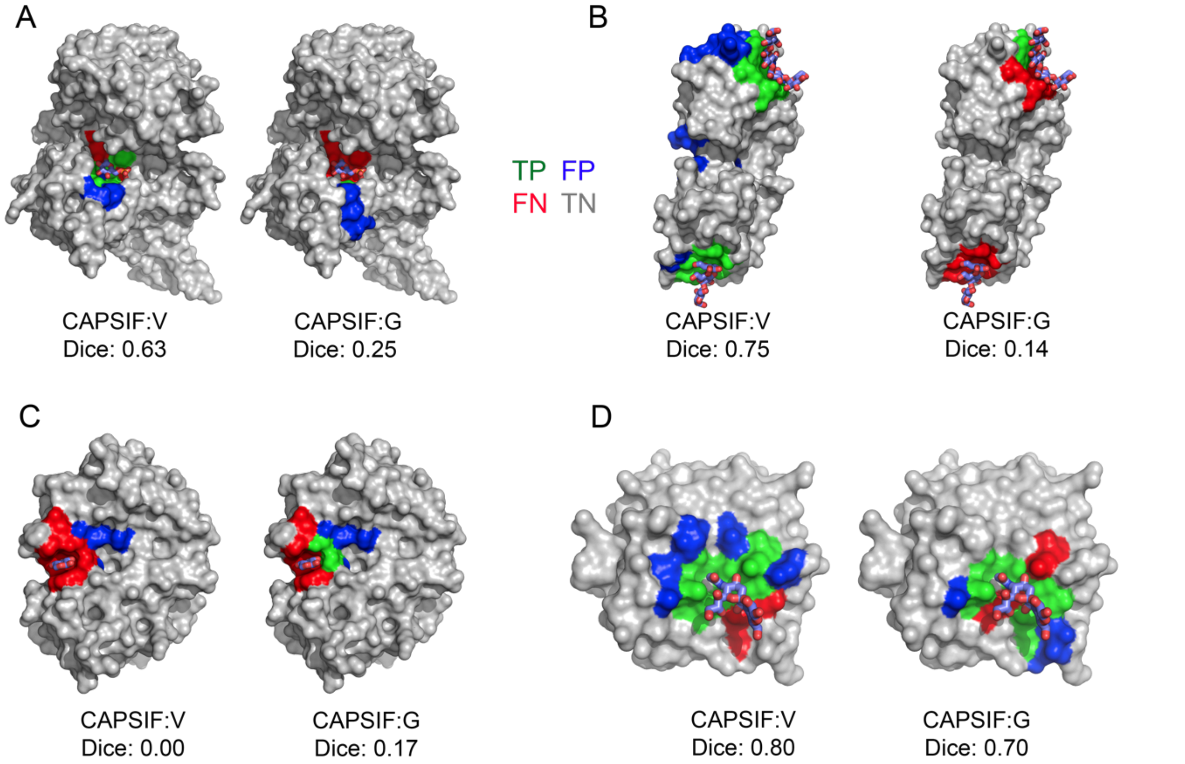


**Figure S6**: **Prediction of carbohydrate binding sites on a protein surface using CAPSIF:V and CAPSIF:G**. **(A)** Glc 6-phosphate dehydrogenase (PDB:5UKW), **(B)** streptococcal virulence factor (PDB:2J44), **(C)** MCR-1 catalytic domain (PDB:5ZJV), and **(D)** CBM40 (PDB:6ER3). Residue labels - green: true positive, blue: false positive, red: false negative, gray: true negative, cyan: bound carbohydrate; Dice coefficient is defined by eq (1) and DCC is distance from center to center of the predicted binding regions.

### Comparison of RCSB and AF2 predicted structures

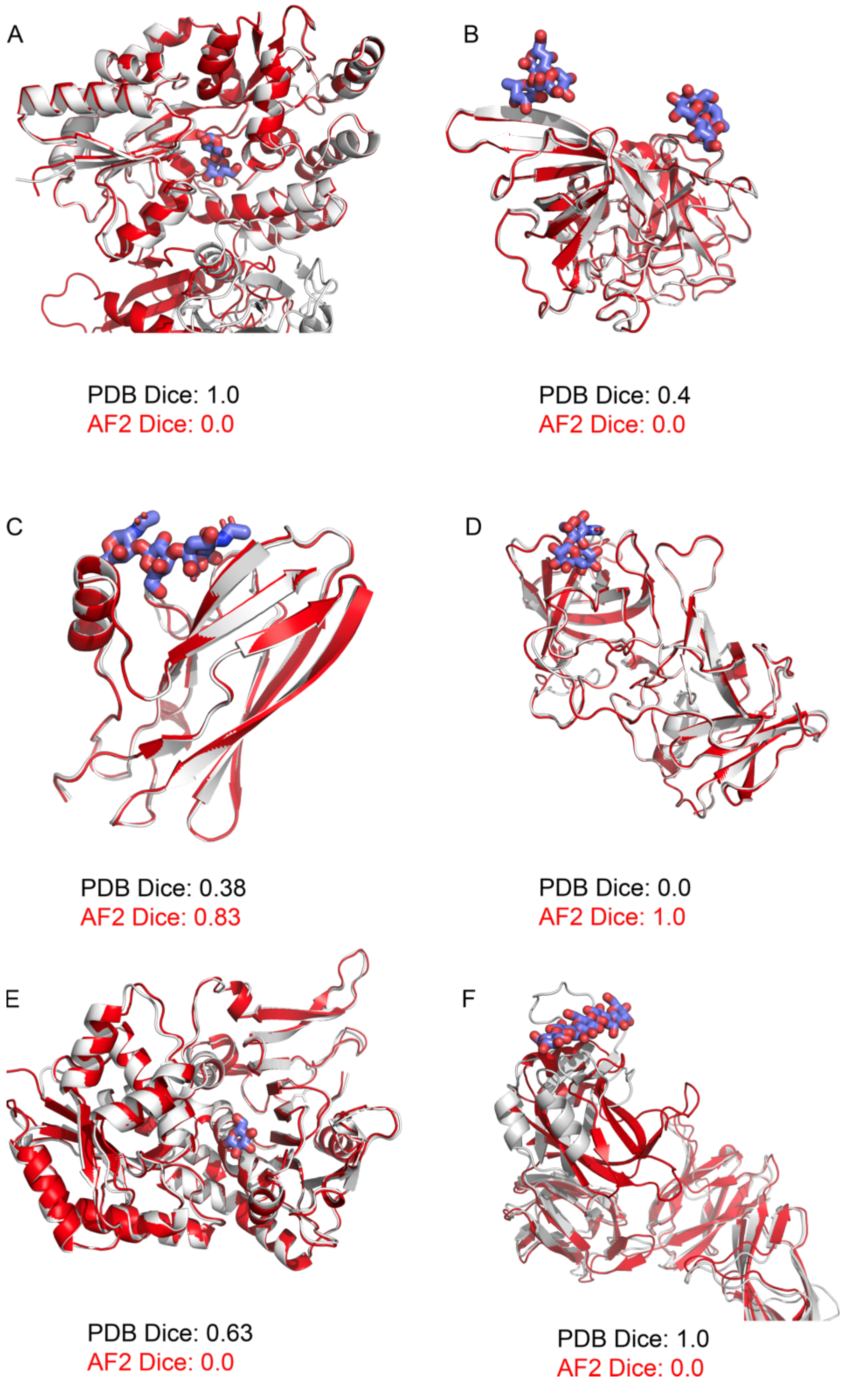


**Figure S7**: **AF2 structure prediction (red) of carbohydrate (purple) binding proteins compared to experimentally solved structures (white)**; **(A)** SUFU (PDB:4BL8) **(B)** *E. coli* aminopeptidase N (PDB:4XO5), **(C)** GspB siglec domain (PDB:5IUC), **(D)** GII.13 novovirus capsid P domain (PDB:5ZVC), **(E)** Glc 6-phosphate dehydrogenase (PDB:5UKW), and **(F)** surface GBP B (PDB:6E57). Dice coefficient is defined by eq (1).

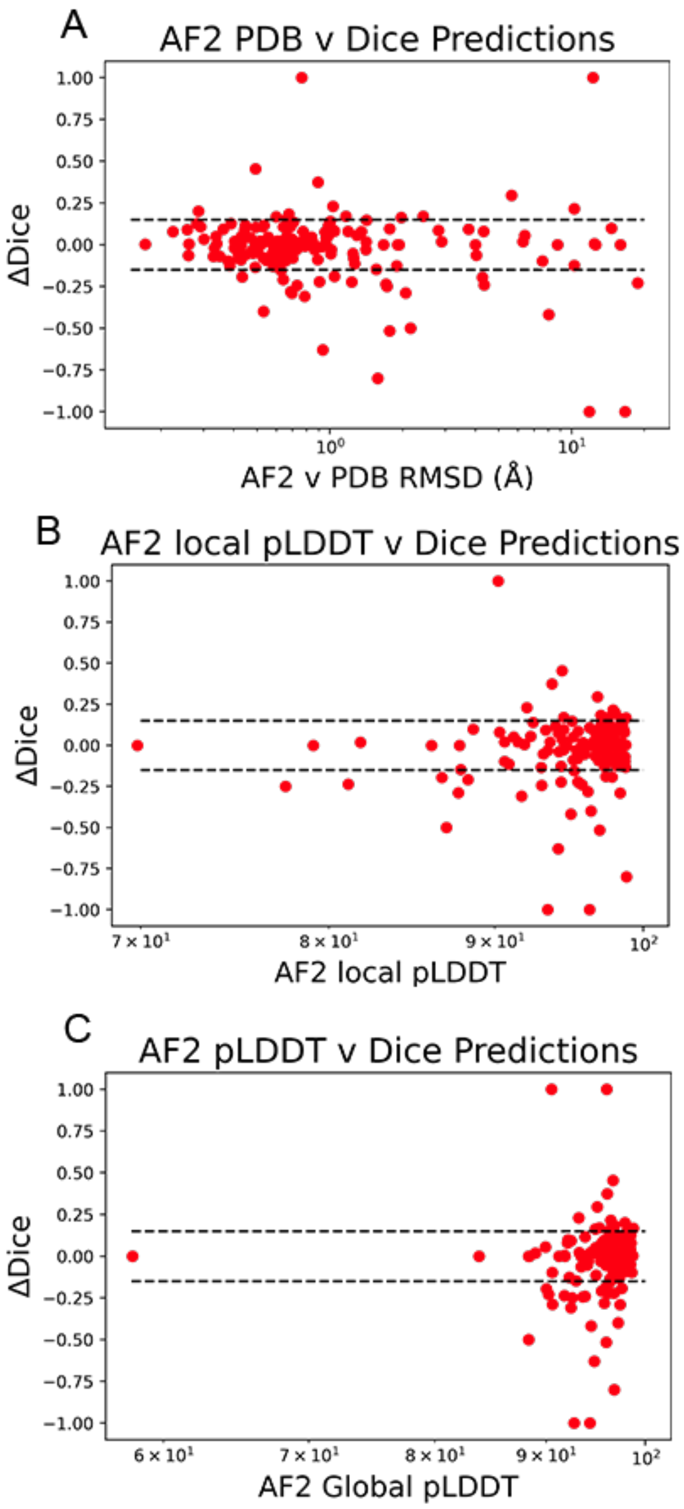


**Figure S8: CAPSIF:V accuracy is not correlated with AF2 accuracy or confidence**. CAPSIF:V predictions on AF2 structure prediction metrics of carbohydrate binding proteins compared to RCSB structures. Change in Dice metric (Δ*Dice* = *AF2 Dice* – *RCSB Dice*) compared to **(A)** the total Cα RMSD (log scale), **(B)** Local average pLDDT score of the carbohydrate binding region, and **(C)** total average pLDDT score of the entire structure.
